# Air Pocket for a Non-Air-Breathing Fish: Developing Respirometry Methodology for the critically endangered Delta Smelt

**DOI:** 10.1101/2024.08.23.609307

**Authors:** Garfield T. Kwan, Florian Mauduit, Kenneth W. Zillig, Yuzo R. Yanagitsuru, Sarah E. Baird, Dennis E. Cocherell, Nann A. Fangue

## Abstract

Respirometry quantifies O_2_ consumption to estimate metabolic activity across stressor(s), and the resulting data is central for devising conservation strategies for fishes. Due to its rapid adoption and widespread utility, protocols have been established to foster quality control and increase consistency. However, the application of established respirometry protocols on the Delta Smelt (Hypomesus transpacificus) have been largely unsuccessful, with reports over the past 12 years documenting high mortality rates that can only be circumvented with very short measurement duration (∼4 h). This prevents the measurement of standard metabolic rate (SMR) and critical O_2_ tension (P_crit_), and ultimately obstructed physiological research, management actions and conservation tools development. In this study, we detailed the anecdotal, behavioral and empirical evidence that culminated in successful measurements of Delta Smelt SMR and P_crit_ at 10, 12, 15, 17 and 19°C. We discovered the Delta Smelt is physotomous and requires frequent refilling of its air bladder to maintain buoyancy. Inspired by air-breathing fish respirometry, the inclusion of an air pocket within their chamber led to considerably more robust respirometry measurements compared to past attempts. The influence of fasting and time of day on O_2_ consumption were also examined. Altogether, this methodology paves the way for acquiring critical knowledge needed to inform Delta Smelt research and conservation (e.g. the development of metabolic index). The lessons learned may also have implications for other aspects of Delta Smelt conservation such as decreasing conservation aquaculture operating costs and improving survival during fish transport and supplementation.

## Introduction

The demand for physiological tools in conservation research is steadily increasing (Fangue et al., 2022). In fish, bioenergetics and respiratory physiology are considered particularly crucial for understanding their ecological performance (Fry, 1971). Environmental temperature significantly influences the physiology and O_2_ requirements of ectothermic fishes. This is further exacerbated by human activities (e.g. damming, eutrophication), which depletes dissolved O_2_ (DO) in aquatic habitats and leads to more frequent episodes of hypoxia (Diaz and Rosenberg, 1995; Breitburg et al., 2009). Consequently, it is argued that a fish’s ability to meet O_2_ demands for essential processes such as swimming, digestion, and assimilation of food may dictate the habitats they can successfully inhabit (Fry, 1971). As a result, traits related to metabolism and energy are increasingly utilized in conservation physiology, both in experimental settings and to inform models projecting the potential impacts of environmental changes and assessing habitat suitability.

Metabolic rate estimation relies on respirometry, a method that measures the gas exchange rates between an organism and its environment (Nelson, 2016). Specifically, the O_2_ uptake rate of an organism is expected to be stoichiometrically related to rates of ATP production through mitochondrial oxidative phosphorylation, thus serving as a proxy for metabolic rate (Nelson, 2016). Respirometry classically allows the measurement of four traits related to metabolism: the standard metabolic rate (SMR), the maximum metabolic rate (MMR), the aerobic Scope for activity (AS), and the critical O_2_ tension (P_crit_). The SMR is the minimum rate of ATP use required to sustain life for a fish that is in a post-absorptive, calm, inactive state after proper thermal acclimation (Chabot et al., 2016a). Being an obligatory expense, SMR is a fundamental and non-negotiable aspect of a fish’s energy budget. All other costs, such as those associated with growth, reproduction, and activity, are additional to the SMR. Using intermittent-flow respirometry, SMR is estimated by collecting measurements of O_2_ uptake over an extended period on an undisturbed animal after acclimation to the respirometry chamber and then calculating a value for SMR using one of a number of statistical methods. The SMR is central to fish energetics as it is an obligatory expense, on top of which all other costs are added (Chabot et al., 2016a). The MMR is the maximum O_2_ consumption rate that a fish can achieve at a given temperature under any ecologically relevant circumstance (Norin and Clark, 2016). Using intermittent-flow respirometry, MMR is determined by exhausting fish either through a chase protocol or a swim tunnel (Rummer et al., 2016; Killen et al., 2017) with MMR taken as the highest rate of O_2_ uptake during recovery (Killen et al., 2021). AS is defined as the maximum capacity to supply O_2_ to sustain metabolic activities beyond SMR and is calculated as the difference between MMR and SMR (Fry, 1971) and is used to predict ecological niche of aquatic ectotherms. By extension, AS can also be used to predict the consequences of future environmental changes using for instance, the debated O_2_- and capacity-limited thermal tolerance theory (OCLTT; Pörtner and Knust, 2007; Clark et al., 2013; Pörtner et al., 2017; Jutfelt et al., 2018). Finally, P_crit_ corresponds to the pO_2_ below which an animal’s basic metabolic needs are not met. Deutsch et al. (2015, 2020) proposed a metabolic index by combining P_crit_ data to climatic, and biogeographic data. This index is used to map the ratio of O_2_ supply to resting metabolic O_2_ demand across geographic ranges of several marine ectotherms and determine suitable ecological niches for the species.

Respirometry protocols have been established to foster quality control and increase consistency of metabolic traits such as SMR, MMR, and P_crit_ (Chabot et al., 2016b, a, c; Killen et al., 2021). This has allowed pursuit of a broad array of research questions from fish behavior (Hansen et al., 2020), ecology (McInturf et al., 2022), aquaculture (Brauner and Richards, 2020), and physiological and climate change responses (Hancock and Place, 2016; Nadler et al., 2016a, 2021; Montgomery et al., 2019; Norin and Metcalfe, 2019; Kwan et al., 2021). However, established standardized respirometry protocols have been largely developed using model species and may not be universally applicable. For example, there is a wealth of research on salmonids (e.g. Eliason et al., 2011; Zillig et al., 2023a, b) which lend themselves well to the established respirometry protocols and equipment. In contrast, there is considerably less respirometry research data on pelagic fishes; of the 115 published respirometry studies aggregated by Killen et al. (2016b), only 18 studies (15.7%) examined pelagic fishes. The lack of respirometry data on pelagic fishes (e.g., Osmeridae, Clupeidae, Scombridae) can be attributed to their aversion to isolation and confined speces, which are inherent to respirometry setups. These complications likely increase their respiration rates and ultimately lead to an overestimation of their MO_2_ (Bernreuther et al., 2013). Therefore, there is a need to develop species-specific respirometry techniques (particularly in pelagic fishes) to enable research on a broader assemblage, especially those of conservation concern.

The Delta Smelt (Hypomesus transpacificus) was one of the most abundant fish species endemic to the San Francisco Estuary (SFE), but their population has declined precipitously - with recent annual surveys regularly reporting few to no detections (Tempel et al., 2021). This catastrophic decline has contributed to their listing as threatened and warranted for endangered status in the United States Federal Endangered Species Act (US Office of the Federal Register, 1993, 2010, 2020), endangered in the California Endangered Species Act (California Fish and Game Commission, 2009), and critically endangered by the International Union for Conservation in Nature Red List of Threatened Species (IUCN, 2014). These listings have spurred research on the species to inform conservation actions and protect the species (Hobbs et al., 2017; Moyle et al., 2018; Tempel et al., 2021; Yanagitsuru et al., 2022; Baerwald et al., 2023). While numerous studies have examined the Delta Smelt’s upper thermal limits (Swanson et al., 2000; Komoroske et al., 2014; Jeffries et al., 2016; Davis et al., 2019), the effect of environmental conditions on their bioenergetic and respiratory physiology remains largely unresolved; in fact, current Delta Smelt bioenergetic models are based on an entirely different species, the rainbow smelt (Osmerus mordax; Lantry and Stewart, 1993; Rose et al., 2013).

Over the past 12 years, adult Delta Smelt respirometry has been largely unsuccessful. The earliest documented attempt at Delta Smelt respirometry was conducted in June 2012, where the use of annular respirometers for 14 hours overnight resulted in 100% mortality (Eder et al., 2013; Table 1). Similarly, the use of glass mason jars for “a few hours” in August 2012 resulted in 100% mortality (Eder et al., 2013; Table 1). At the time, Delta Smelt mortality was attributed to high flow rate, so smaller recirculating pumps and custom-made cylindrical chambers with barriers were used to slow flow rate and generate laminar flow conditions. Testing with these chambers in October 2012 slightly decreased mortality to 91.3% (Eder et al., 2013; Table 1). Later in November 2012, researchers discovered the time of day significantly impacts respirometry success. This was most evident when Delta Smelt were tested at 15°C: overnight respirometry trials that started in the afternoon resulted in 68 – 85% mortality, whereas respirometry that started in the late morning (10am) to noon resulted in 16.7 – 33.3% mortality (Eder et al., 2013; Table 1). Despite this improvement, daytime respirometry of smelt acclimated to 19°C Delta Smelt resulted in 54.2% mortality. Moreover, Delta Smelt frequently exhibited a nosing behavior towards the surface of the chamber, which prompted the design from a circular chamber to a rectangular chamber. To circumvent high mortality rates, respirometry trials were shortened to ∼4 hours, which subsequently reduced mortality to 0 – 5.6% (Table 1). All of these adjustments led to the first publication of Delta Smelt respirometry data reported as routine metabolic rate (Hammock et al., 2017), though the relatively short 4-hour (7 respirometry cycles) duration was likely incompatible with the determination of SMR (Chabot et al., 2016a).

**Table 1:** Past experimental designs on Delta Smelt respirometry.

| Citation | Chamber Name | Date | Fish length (mm) | Fish weight (g) | Fish age (dph) | Chamber Volume (mL) | Visible Consppecific, Ceiling | Lighting | Surface Access | Duration (Hours) | Mortality (%) | Starting Time |
| --- | --- | --- | --- | --- | --- | --- | --- | --- | --- | --- | --- | --- |
| *unpublished data* | Annular respirometer | Jun 2012 | NA | NA | NA | 10,000 | Y, N | No light; covered in plastic tarp | N | 14 | 100.0% | Afternoon |
| *unpublished data* | Glass mason jars | Aug 2012 | NA | NA | NA | 2000 | Y, N | No light; covered in plastic tarp | N | "a few hours" | 100.0% | Afternoon |
| **Eder et al., 2013** | Cylinder chamber | Oct 2012 | NA | NA | NA | 273 - 975 | Y, N | No light; covered in plastic tarp | N | NA | 91.30% | Afternoon |
| **Eder et al., 2013** | Cylinder chamber with flat plate | Nov 2012 | NA | NA | NA | 273 - 975 | Y, N | No light; covered in plastic tarp | N | NA | 68 - 85% | Afternoon |
| **Eder et al., 2013** | Cylinder chamber with flat plate | Dec 2012 | 43 - 62 | 0.6 - 2.1 | NA | 273 - 975 | Y, N | No light; covered in plastic tarp | N | 6 | 11.5 - 54.2% | Mid-morning to noon |
| *unpublished data* | Rectangular chamber | Sep 2013 | juvenile: 50 - 54; adult: 76 - 77 | juvenile: 1.0 - 1.2; adult: 3.2 - 3.7 | NA | juvenile: 250; adult: 1,050 | Y, N | Dim white fluorescent light; covered in plastic tarp | N | 4 | 5.6% | Mid-morning to noon |
| **Hammock et al., 2017** | Rectangular chamber | Nov 2013 | NA | 0.9 - 3.3 | ~221 | 1,250 | Y, Y | Dim white fluorescent light; covered in plastic tarp | N | 4 | 0% | Mid-morning to noon |
| **This study** | Rectangular chamber with surface access | Oct 2023 | 54 - 84 | 1.0 - 5.2 | 259-302 | SMR: 500; P<sub>crit</sub>: 700 | Y, Y | Overhead red light | Y | 17 - 21.5 | 40% | Afternoon |
| **This study** | Rectangular chamber with surface access | Oct and Nov 2023 | 54 - 84 | 1.0 - 5.2 | 259-302 | SMR: 500; P<sub>crit</sub>: 700 | Y, Y | Overhead red light | Y | 10 - 12 | 5% | Before dawn |

The objective of this study was to determine the effect of temperature on the bioenergetic and respiratory physiology of adult Delta Smelt. Specifically, we aimed to improve Delta Smelt respirometry methodology through the incorporation of red lighting and surface access to more accurately measure their SMR and P_crit_. In addition, we tested the influence of time of day (daytime vs overnight) and fasting on the organism’s O_2_ consumption across a broad range of ecologically relevant acclimation temperatures including 10, 12, 15, 17 and 19°C.

## Result and Discussion

### Lighting and Delta Smelt handling

SMR measurements of species active during the day (e.g. salmon) is typically conducted in near to complete darkness in an effort to calm the fish, and to prevent sudden changes in lighting to influence respirometry measurements (Speers-Roesch et al., 2018). However, because Delta Smelt are a shoaling fish species, the inability to see conspecifics is hypothesized to induce stress. Indeed, past studies in other shoaling species have reported that visual cues of conspecifics can influence MO_2_ (Hall and Clark, 2016; Killen et al., 2016b; Nadler et al., 2016b). Therefore, initial prototypes of Delta Smelt respirometry chambers were designed with clear walls and placed in close proximity to allow vision of conspecifics. However, past Delta Smelt respirometry trials blocked ambient lighting from entering the experimental area with black plastic tarps (Table 1). Given that Delta Smelt collisions with fish exclusion screens is more frequent in dark conditions (Swanson et al., 2005), the lack of lighting during past Delta Smelt respirometry trials may similarly impeded vision of conspecifics and increased collisions within the chamber, and ultimately contributed to greater MO_2_ and increased mortality (Table 1).

The rearing protocols of the Delta Smelt refuge population at the Fish Conservation and Culture Laboratory (FCCL, Byron, CA, USA) suggest this critically-endangered fish is very light-sensitive. Past results have documented indoor lighting promotes hatching and feeding at the early larval stage (<40 days post hatching (dph)), but by the late larval stage (40-80 dph) Delta Smelt prefer lower light/higher turbidity conditions (Tigan et al., 2020). This continues until maturity (∼300 dph), when they are transferred to outdoor tanks and that dampens ambient light with shade cloth tank lids to promote sexual maturation and spawning (Hung TC, *personal communication*). Current best practices in Delta Smelt transport and rearing require carboys and tanks to be painted black to reduce overhead shading (Cocherell DE, personal communication).

Despite the extensive respirometry guidelines compiled by Killen *et al*. (2021), lighting is rarely reported as it is (or assumed to be) generally conducted in the dark or heavily shaded to block indoor fluorescent lighting. Of the full light spectrum, red light is known to dissipate rapidly as it permeates through the water column and thus thought to be less disruptive to fishes. While it is less disruptive than white light, there is some evidence even deep-sea fishes can still perceive red light (Widder et al., 2005). The impacts of different colored lighting are species-specific (Ruchin, 2020), and have been shown to impact an array of physiological parameters including feeding and growth rate (Ruchin, 2004; Volpato et al., 2013), cortisol levels (Owen et al., 2010), liver and gonad conditions (Yuan et al., 2017).

Red lighting may have other biological effects: in one study on the Nile tilapia (*Oreochromis niloticus*), red light induced greater feeding behavior though it intriguingly did not significantly increase growth rate (Volpato et al., 2013). In this study, Delta Smelt under red lighting (9 – 31 lumens) were observed to remain calm and not react to overhead shadows, yet able to see conspecifics and align themselves accordingly (Supplemental video 1) as well as form and maintain shoals (Kwan, personal observation).

The sole illumination with red lighting can also be thought of as the omission of other wavelengths, which perhaps simulates turbidity for the Delta Smelt. In the wild, Delta Smelt abundance is closely associated with an optimal water turbidity (Hamilton and Murphy, 2022; Smith and Nobriga, 2023) and linked with increased feeding efficiency and reduced predation risk (Ferrari et al., 2014; Mahardja et al., 2016; Schreier et al., 2016; reviewed in Yanagitsuru et al., 2022). In this study the use of red light allowed us to observe respirometry trials with minimal disturbance to this sensitive fish, and to conduct the experiment more safely and efficiently. Though their shoaling behavior suggests the Delta Smelt is able to see under the red lighting, future research should determine their spectral sensitivity to determine their visual capacities. Moreover, future studies should seek to compare and quantify their metabolic rate and stress levels across different light wavelength, lighting intensity and turbidity levels to and ultimately verify whether red lighting can be a useful substitute for turbidity. If future research reveals red lighting decreases Delta Smelt metabolic rate and stress levels, then conservation hatcheries such as the FCCL could employ these lighting strategies to reduce operational costs associated with the use of turbidity and increase safety and efficiency of animal husbandry routines as proposed for freshwater catfish (*Wallago attu;* Giri et al., 2002).

### Surface access is necessary for buoyancy, survival

The prolonged lack of surface access during previous transport and field release attempts may explain high Delta Smelt mortality. Initially, plastic buckets (∼19 L) were fitted with seine to prevent fish from escaping during short-distance transport, but Delta Smelt were often caught in the mesh – leading to struggling and mortality (Cocherell, D. *personal communication*). At the time, this behavior was interpreted as an attempt to escape, so the Delta Smelt transport protocol was altered to using flat plastic lids to remove air pockets. Later iterations required transport over longer distances, and the standard operating procedures of Delta Smelt transport within insulated carboys (∼83 L) continued to eliminate the air pocket to prevent injury and mortality caused by splashing (Cocherell, D. *personal communication*). Transport mortality was further reduced with NaCl addition (∼5 ppt) (Swanson et al., 1996); the higher salinity provided greater environmental counterions to facilitate internal homeostasis maintenance throughout transport, during which CO_2_ accumulates, water pH decreases (Hudson et al., *in prep*), and nitrogenous waste is accrued.

Delta smelt appears to be able to tolerate ∼4 hours of carboy transport without surface access with minimal mortality (<1%). In contrast, transport of ∼5 hours had resulted in ∼6.25% mortality (∼200 of 3,200 fish) (Dennis personal comm?). Later, as conservation focus shifted towards the release of hatchery raised Delta Smelt to supplement wild populations, researchers tested field cages to allow the fish to acclimatize to the wild prior to their release. Since Delta Smelt were thought to be poor swimmers (Swanson et al., 1998, 2000), early prototypes sought to reduce wave action by submerging the field cages. Similar to carboy transport reports, underwater submergence led to very high mortality (Cocherell, D. *personal communication*), whereas later iterations that buoyed the field cages to the surface greatly reduced mortality to <1% (USFWS, 2022, 2023).

The use of red lighting allowed for extensive observation of Delta Smelt behavior throughout respirometry trials, which could not be done with past methodologies as they were performed in dim lighting or complete darkness. During initial troubleshooting, Delta Smelts were submerged within cylindrical respirometry chambers (volume = ∼500 cm^2^) for 4 – 12 hours, and bubbles have been observed to pop out of the mouth of the fish (Kwan GT, *personal observations*). Upon release, Delta Smelts would doggedly and frequently swim to the surface to air gulp despite normoxic conditions in both the respirometry chamber and the holding tank (Kwan GT, *personal observations*). Of the fish that died during respirometry trial, bubbles were observed within the chamber. Post-mortem autopsy revealed Delta Smelt are physotomous (air bladders are connected to their esophagus), and comparison to healthy fish in holding tanks revealed submerged fish exhibited a reduced air bladder volume (Figure 1). All of these observations imply surface access restriction led to air bladder deflation.

**Figure 1.**
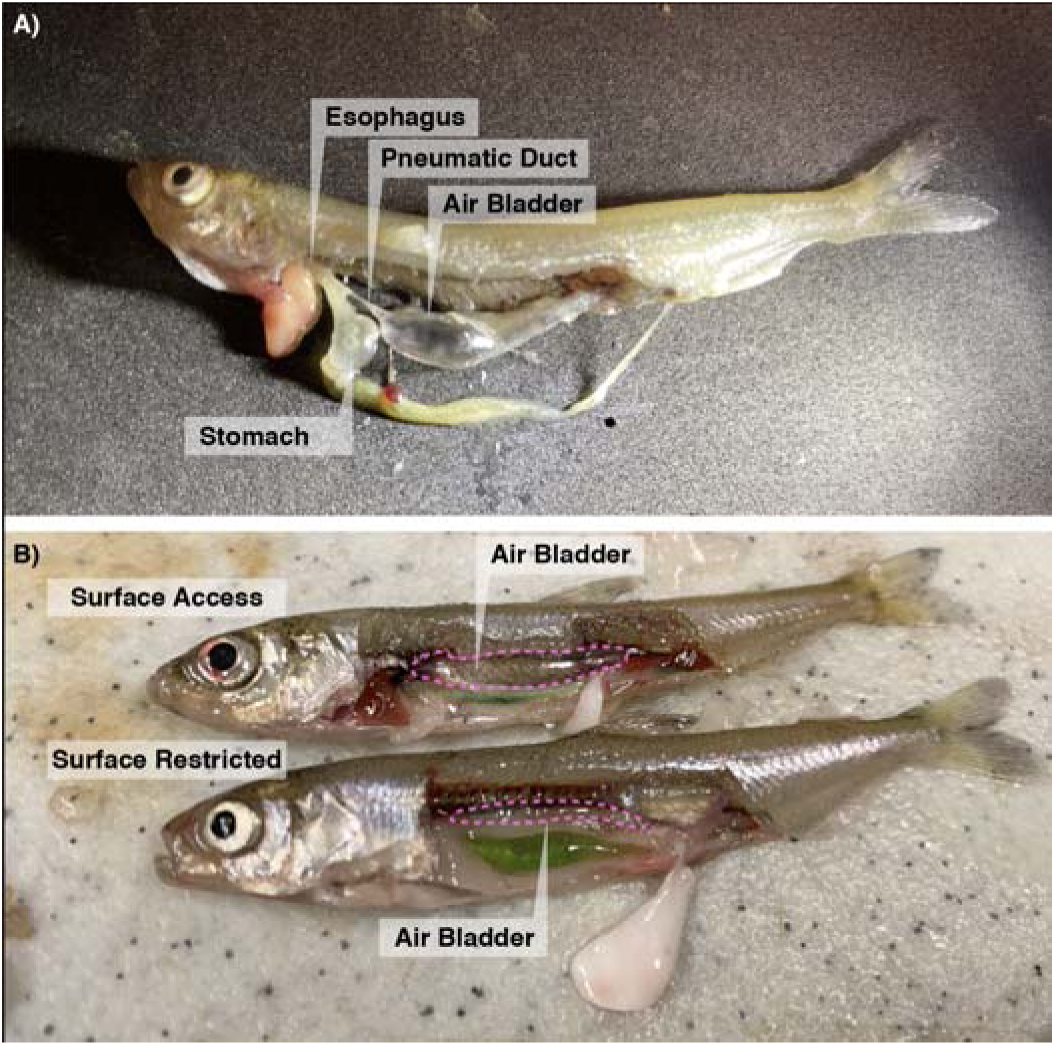
Delta Smelt is physotomous. **A)** The esophagus of the Delta Smelt connects to the stomach, and to the air bladder through the pneumatic duct. **B)** Air bladder appears to be deflated after 8 hours-long respirometry without surface access. Pink dotted lines indicate the perimeter of the air bladder.

In retrospect, Delta Smelt experienced surface access restriction during tank and carboy transport, submerged field cage deployment, and within the various iterations of respirometry chambers. Delta Smelt attempts to reach the surface to refill their air bladder led to entanglement, higher swimming activity and energy expenditure, and eventually exhausting and mortality despite normoxia within respirometry chambers. Subsequent observations during field release further bolstered our findings: Delta Smelt released into field cages were witnessed to repeatedly air gulp following carboy transport (Cocherell D, *personal observations*).

Curiously, other fishes commonly used in respirometry experiments are also physostomous. For instance, Chinook salmon (*Oncorhynchus tshawytscha*) have survived enclosure within respirometry chambers for ∼5 days (Lo et al., 2022). Even so, salmonids eventually require surface access: Atlantic salmon (*Salmo salar*) are known to experience buoyancy problems after three weeks of submergence (Dempster et al., 2009; Korsøen et al., 2009). Why Delta Smelt lose air bladder volume more rapidly than other physotomous fishes is not known, but this characteristic is reflected in their adaptation to their historical habitat and emphasizes their vulnerability under heavily altered current-day conditions. Historically, Delta Smelt habitats were shallow and muddy, whereas current-day conditions are relatively deeper and significantly less turbid (Moyle et al., 2016). This implies Delta Smelts are negatively affected by habitat compression as the deep channelized rivers provide less surface access than shallow floodplains. Given that the tendency of gases escaping from within the air bladder is proportional to water depth (Strand et al., 2005), Delta Smelt inhabiting today’s channelized river conditions may be losing their buoyancy at a greater rate if they reside deeper in the water column during the day, and may thereby become more vulnerable to predation as they more frequently access the surface to refill their air bladder. Altogether, the Delta Smelt’s close association with the surface further strengthens the necessity to restore historical floodplain habitats around the SFE in order to help their populations recover.

When housed in groups of 2 to 4 in a larger tank (volume = ∼3,400 cm^2^), most (75%; 6 of 8 fish) Delta Smelt can survive 4.5 days without surface access (Figure 2). As the duration of surface restriction increased, Delta Smelt buoyancy became increasingly negative (Figure 2) and must keep swimming while angled towards the surface to avoid sinking. Delta Smelts that were denied surface access were observed to rapidly swim upwards or rest on the bottom. However, this predicament was reversible: Delta Smelt recovered their buoyancy when given surface access over 2.5 days (Figure 2I, J; Supplemental Video 2). Altogether, this suggests earlier Delta Smelt respirometry measurements without surface access likely resulted in elevated SMR levels as the fish must expend greater energetics to maintain their buoyancy or in their attempt to reach the surface.

**Figure 2.**
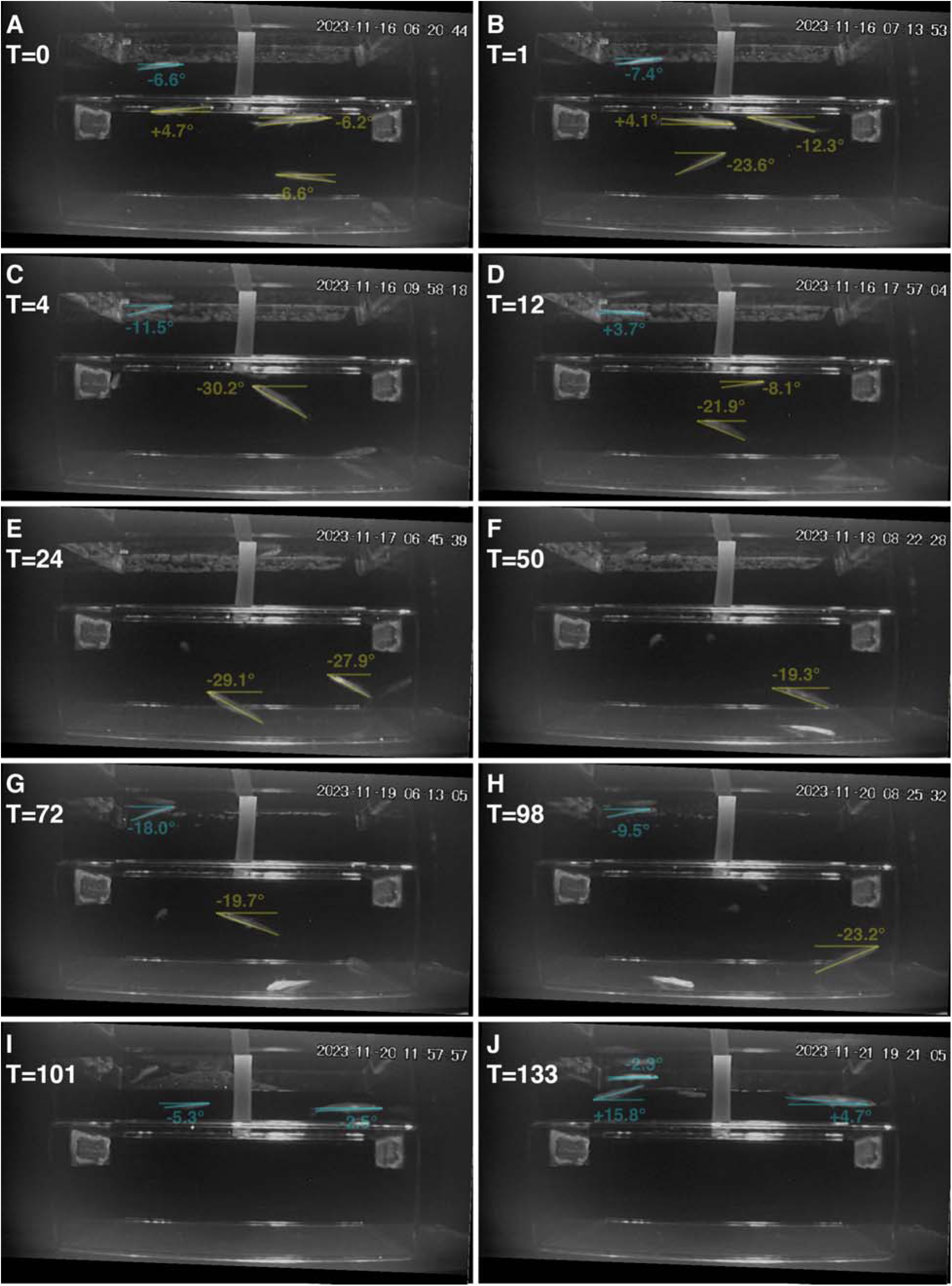
Video screenshots of Delta Smelt during forced submergence and subsequent recovery. Delta Smelt after A) 0 hour, B) 1 hour, C) 4 hour, D) 12 hour, E) 24 hour, F) 50 hour, G) 72 hour, and H) 98 hours without surface access. After 100 hours, Delta Smelts were allowed surface access, and their buoyancy recovered rapidly I) within 1 hour (T=101) and maintained J) after 33 hours (T=133). One fish died after 48 hours without air access. The angles of Delta Smelt snout to caudal peduncle were measured, with blue lines representing fish with surface access whereas yellow lines represent fish without surface access. Images were adjusted by 2.2° due to a tilt in the camera prior to measurements. Time stamp on the top right is denoted as year-month-day hour-minute-seconds.

### Incorporating surface access into respirometry

This study is the first to recognize that Delta Smelt require surface access to maintain their air bladder volume, and its incorporation into respirometry chambers have allowed Delta Smelt to survive significantly longer durations compared to past iterations (Table 1). In general, respirometry setups take great pain to remove air bubbles from the chamber as O_2_ diffusion from the air into the water could skew measurements. In order to reconcile the Delta Smelt’s need for surface access, we drew upon past bimodal respirometry chamber designs used for air-breathing fishes (Lefevre et al., 2016). In this study, air pockets were created within the Delta Smelt respirometry chambers by positioning individual chambers slightly above the water line (Figure 3). This allows the water and air levels within the chamber to equilibrate with the water height of the external water bath. Under red lighting, Delta Smelt were observed to utilize the provided air pocket within the respirometry chamber (Kwan, *personal observation*).

**Figure 3.**
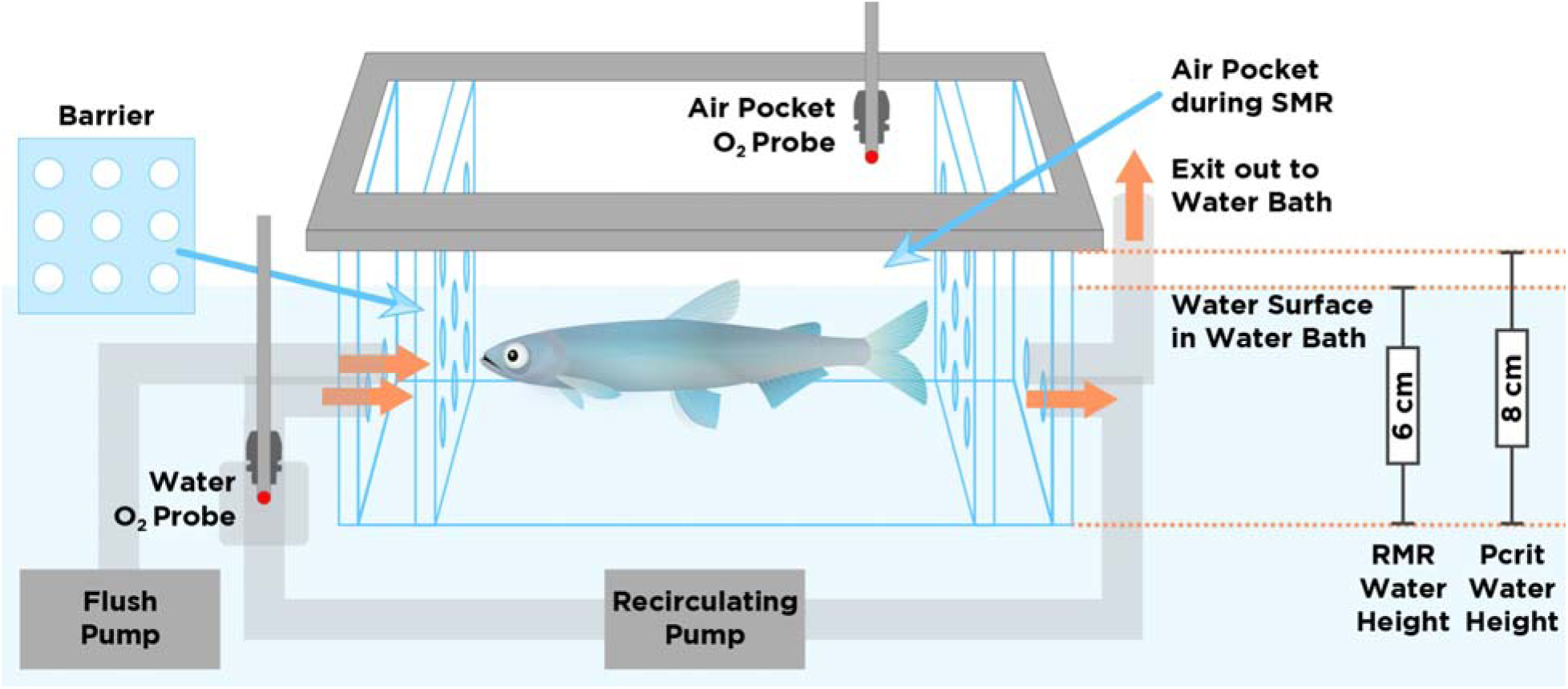
Schematic of Delta Smelt respirometry setup. Delta Smelt respirometry was conducted in a rectangular acrylic apparatus, and two optical fiber probes simultaneously measured O_2_ in both water and air pocket. Two barriers allowed water flow but prevented the fish from swimming into the tubing. During standard metabolic rate (SMR) measurements, an air pocket provided Delta Smelt with surface access. To prevent air intake, the recirculating pump and flush pump intakes were placed near the bottom of the chamber. Air pocket was removed during critical O_2_ tension (P_crit_) analysis by water flushing and the temporary removal of the air O_2_ probe. See the methods section below for additional details.

One major concern was that the air pocket could diffuse O_2_ into the water, which would artificially reduce MO_2_ during respirometry trials as O_2_ consumed by the fish is replaced by O_2_ from the air pocket. To assuage these concerns, we simultaneously measured O_2_ in both the water and the air pocket throughout the respirometry trials, then measured the diffusion rate of the air pocket and compared it to Delta Smelt metabolic rates. On average, Delta Smelt typically consumed ∼1-3% of the DO over the 13-minute respirometry cycle. To determine the effects of the air pocket on DO at 19°C (the highest temperature tested), we set up an empty respirometry chamber with 97% DO (without fish) and measured water and air O_2_ over a 3-hour period. Rather than an increase in water DO indicative of greater air to water diffusion, we observed a slight decrease throughout the 3-hour period. To compare, the average Delta Smelt SMR at 19°C consumed O_2_ at a rate of 243.1 ± 26.5 mg O_2_ kg^-1^hr^-1^ (mean ± SEM; n=14), whereas the no fish control recorded a decrease at a fish-standardized rate of 6.6 ± 1.6 mg O_2_ kg^-1^ hr^-1^ (mean ± SEM; n=4) or ∼2.7% of fish MO_2_ at 19°C. This suggests any contribution by air-water diffusion is negligible, and likely masked by the presence of background respiration. However, when DO in the water was further reduced as typical of P_crit_ trials, O_2_ diffusion from the air pocket into the water became evident. Despite Delta Smelt respiration, our early troubleshooting trials detected an increase in water DO and a positive MO_2_ slope when nitrogen bubbling brought DO levels below 80%. This suggests the air pocket had negligible impact during SMR measurement (when water DO is cycled between 97-100%) but that it must be removed during stepwise O_2_ drawdown during P_crit_ measurement. As such, our Delta Smelt respirometry protocol details the removal of the air pocket when we transitioned from SMR to P_crit_ trials (Figure 3; see Methods below).

### Influence of time of day

The inclusion of an air pocket successfully reduced overnight respirometry mortality from 100% to 40% (Table 1), and generated the longest Delta Smelt respirometry measurements to date. While this was a significant improvement, we desired further reduction in mortality levels. Since previous attempts had documented considerably higher survival when respirometry measurements were collected during the daytime (Table 1), we decided to switch to starting before dawn (predawn) and measured MO_2_ throughout the day. This change further reduced Delta Smelt mortality from 40% to 5%, though this increase in survival could also be related to the shorter duration within the respirometry chamber; overnight trials lasted 17-21.5 hours whereas predawn trials were 10-12 hours in duration. As such, it was necessary to determine whether the shorter predawn trials would yield comparable SMR and P_crit_ measurements as the longer overnight trials.

In this study, we found predawn and overnight techniques generated similar SMR measurements in both 10 and 19°C Delta Smelt (p=0.9832; Figure 4A). As expected, we also detected a significant temperature effect (p=0.0005) with fish tested at 19°C having higher SMR, and their interaction was not significant (p=0.9882; Figure 4A). In contrast, the P_crit_ of 19°C fish estimated with the overnight method was significantly higher than the 19°C fish estimated with the predawn method (p<0.0001; Figure 4B). P_crit_ was also affected by temperature (p<0.0001) but not their interaction (p=0.1579; Figure 4B). We suspected the unnaturally high P_crit_ (e.g. ∼90% DO) were derived from moribund fish, so shorter respirometry durations would likely yield more accurate measurements at less optimal temperatures. However, SMR estimations did not encounter this issue as this value is calculated from the mean of the lowest normal distribution (see methods below), which reflects the period after the Delta Smelt had settled down. Altogether, the shorter 10 to 12 hour-long respirometry trials incurred lower mortality, generated comparable Delta Smelt SMR as the overnight trials, and provided a more accurate P_crit_ estimate.

**Figure 4.**
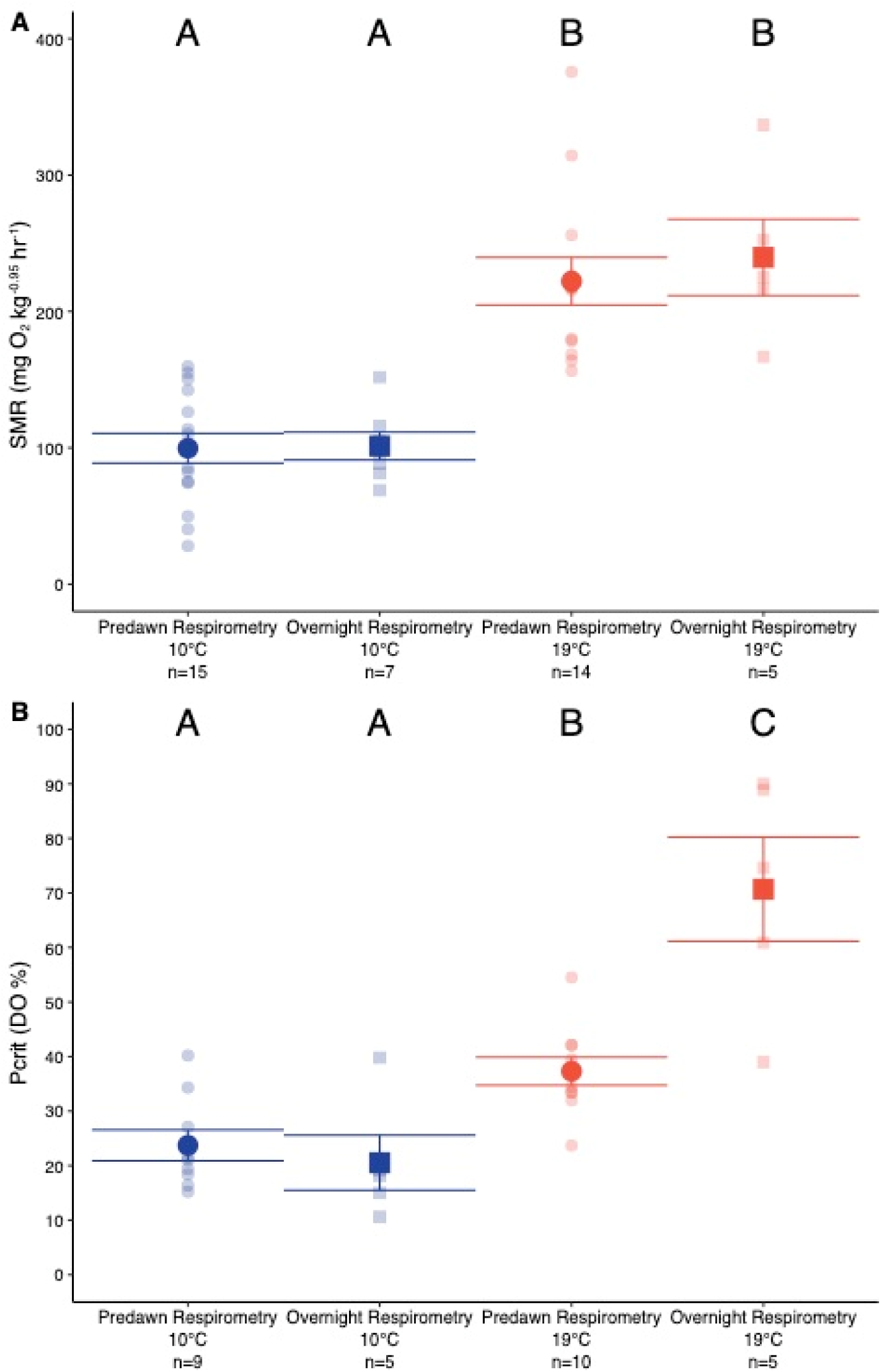
Predawn and overnight respirometry comparison. The starting time and duration in the respirometry chamber did not significantly influence **A)** standard metabolic rate (SMR) of 10°C and 19°C-acclimated Delta Smelt. **B)** Critical O_2_ level (P_crit_) in 10°C-acclimated Delta Smelt was not affected by starting time and duration, whereas P_crit_ was significantly higher in 19°C-acclimated fish that underwent overnight respirometry trials. Data is shown as mean ± SEM. Alphabetic letter denotes significant difference (α=0.05).

Circadian rhythm patterns are known to influence respirometry measurements (Livingston, 1971; Kim et al., 1997; Svendsen et al., 2014), and past studies have indicated Delta Smelt prefer to feed (Hobbs et al., 2006), spawn (Tsai et al., 2022) and shoal with conspecifics (Chase et al., 2023) during the daytime. In the wild, Delta Smelt have also been observed to intentionally change their position within the water column according to the tides (Feyrer et al., 2013). Altogether, the elevated mortality implies the Delta Smelt do not prefer being near the surface during the night, and may have exhausted themselves as they struggled to swim deeper into the water column. As such, the greater success with daytime Delta Smelt respirometry despite similar chamber designs (Table 1) is not entirely surprising. Future studies should examine the interplay among time of day and water column position, and perhaps design and test a respirometry chamber integrated with greater vertical space.

### Influence of fasting

Standard respirometry practices advise fasting fish prior to SMR and P_crit_ trials since feeding is known to increase metabolic rate, thereby inducing greater variance due to differences in food ingested and energetic costs associated with digestion (Chabot et al., 2016c). However, fasting fish prior to respirometry may not always be an option, especially when working with endangered species and at numbers limited by federal/state permits. As such, we examined the impacts of feeding and 2-day fasting on SMR and P_crit_ levels in 10 and 12°C-acclimated Delta Smelt. Both SMR and P_crit_ were not significantly affected by fasting (SMR: p=0.3247; P_crit_: p=0.1811) nor were their interaction with temperature impacted (SMR: p=0.3948; P_crit_: p=0.1983; Figure 5). In contrast, SMR (p=0.0446) but not P_crit_ (p=0.8616) was affected by acclimation temperature (Figure 5). While we acknowledge that our low sample size could have skewed our results, these findings are surprising as feeding typically increases metabolic rates (Goodrich et al., 2022; Lo et al., 2022). In a previous food consumption study, both juvenile and adult Delta Smelt reared at 10 to 18°C completed digestion within 20 hours of *ab libitum* feeding (Eder et al., 2013). Although we did not verify the stomach contents of our fish, this suggests it was unlikely that digestion interfered with the respirometry measurements. Instead, the trend for increased SMR after the 2-day fasting could reflect increased foraging activity given these hatchery-reared fish have been habituated to daily feeding throughout their lives. Perhaps the differences between fed and fasted fish would be more apparent after a longer fasting periods and at higher temperatures, and should be confirmed in future studies.

**Figure 5.**
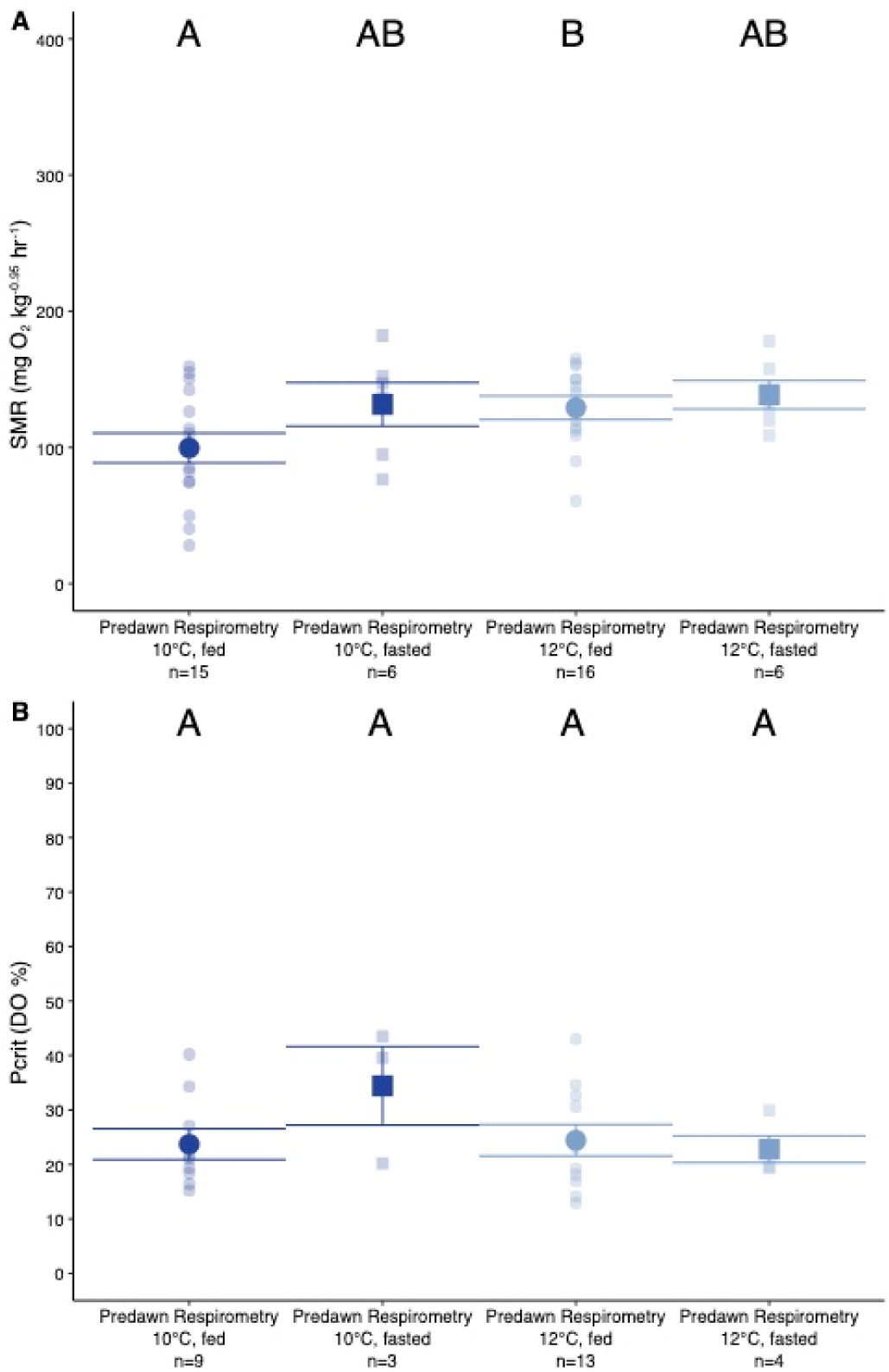
Fed and fasting comparisons. Fasting did not significantly influence **A)** standard metabolic rate (SMR) nor **B)** critical O_2_ level (P_crit_) of 10 and 12°C-acclimated Delta Smelt. Data is shown as mean ± SEM. Alphabetic letter denotes significant difference (α=0.05).

### Comparison to past respirometry data

Altogether, this study documents our extensive efforts over the last 12 years and highlights the recent breakthroughs in accurately estimating physiologically relevant SMR and P_crit_, marking a significant milestone in the determination of Delta Smelt physiological performance and environmental tolerance. Our progress is most evident when compared to two past juvenile and adult Delta Smelt respirometry datasets (Figure 6). The unpublished dataset was collected from juvenile and adult Delta Smelt in September 2013 over a 4-hour measurement period (four 51-minute measurement cycles) under low lighting and with no surface access (Table 1). In another study, Hammock *et al*. (2017) measured 16°C-acclimated fish under low lighting, with no surface access, and limited to 4 hours in duration (five 30-minute measurement cycles) to limit mortality (Table 1). The study ascribed their respirometry measurement as mass-specific resting metabolic rate, and they found salinity (0.4 – 12 ppt) to not significantly MO_2_ (Hammock et al., 2017). Despite a substantially greater duration within the respirometry chamber (Table 1), our inclusion of red light and air surface access resulted in significantly lower SMRs in both 15°C-(p=0.0008) and 17°C-acclimated fish (p=0.0038) compared to Hammock et al. (2017) (Figure 6). Therefore, the MO_2_ measurements presented in Hammock *et al*. (2017) is likely an overestimation as the lack of air surface access may have resulted in increased swimming activity, and the relatively few measurement cycles likely did not overlap with the Delta Smelt’s resting state.

**Figure 6.**
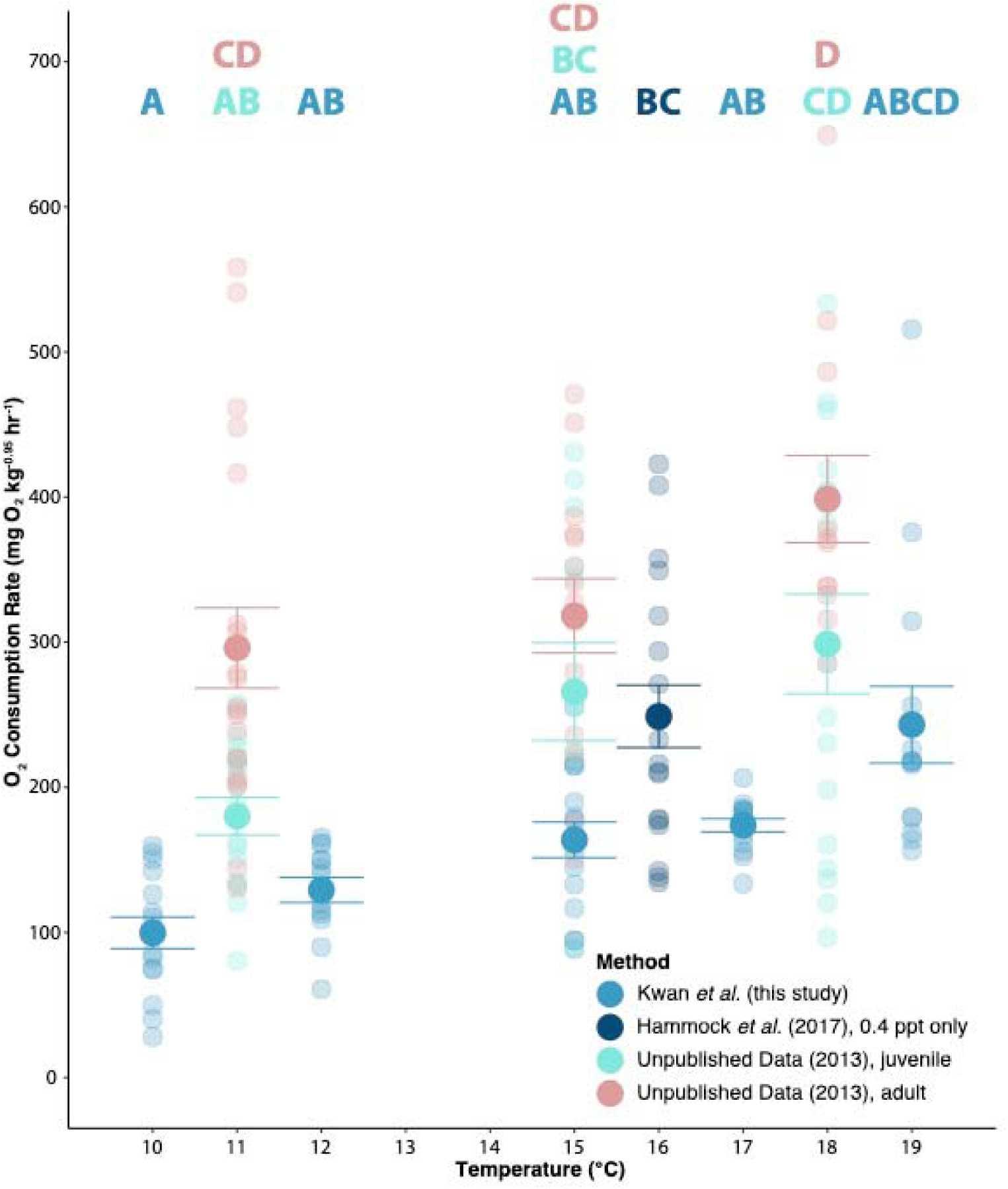
Delta Smelt respirometry comparison. Comparison of respirometry measurement of O_2_ consumption rate across methodologies, including unpublished data collected in 2013, data from Hammock *et al*. (2017), and those described in this study. Data is shown as mean ± SEM. Different alphabet letters denote significant difference (α=0.05).

### Implications and Future Directions

This study summarized the 12-year effort to optimize Delta Smelt respirometry, which culminated in the incorporation of an air pocket for successful SMR and P_crit_ measurements and ultimately unlocked a new frontier in conservation physiology. Respirometry on pelagic fishes are rare (Killen et al., 2016a) due to their unique demand for space, schooling or shoaling behavior, and (in the case of Delta Smelt) air surface access. It is our hope that the present study will inspire further troubleshooting attempts in order to continue filling this gap in knowledge for pelagic fishes.

Besides its utility in generating ecophysiology measurements, these findings can also optimize existing aquaculture rearing, transportation and experimental protocols, and conservation management of this endangered pelagic fish. For instance, research to deduce whether Delta Smelt perceives the omission of certain wavelengths to be similar to turbidity could facilitate animal husbandry and lower operating costs. Next, the determination of the rate at which Delta Smelt loses (and recovers) their buoyancy could bolster their survival during long-term transport. Furthermore, swim tunnel measurements could potentially benefit from the incorporation of an air pocket as in the present study. Moreover, respirometry allows for the quantification of sublethal stressors. For instance, tagging Delta Smelt has been of great interest to conservation managers, but the post-surgical impacts can be sublethal and not apparent when assessed with survival alone. Instead, analyzing post-surgical impacts with SMR measurements could be a more useful in determining the least invasive procedure, and whether a Delta Smelt has recovered from tagging effects. Finally, conservation managers can reliably utilize respirometry to estimate energy consumption across abiotic and biotic stress(ors), and makes feasible the modeling of a Delta Smelt-specific metabolic index.

The modeling of the metabolically viable habitats of Delta Smelt across space and time will be a valuable tool for conservation managers, but it could be further improved if turbidity was incorporated. The relationship between turbidity and Delta Smelt success has been well documented, and higher turbidity has been shown to promote feeding success, growth rate, survival (Hasenbein et al., 2013; Tigan et al., 2020) and reduced cortisol levels in larval (Hasenbein et al., 2016), juvenile and adult Delta Smelt (Lewis et al., 2021; Pasparakis et al., 2023). Conversely, reduced turbidity across the SFE has been correlated with higher cortisol level (Pasparakis et al., 2023) and increased predation risk (Moyle et al., 2016). Since Delta Smelt must frequently access the surface to maintain nominal buoyancy, low turbidity conditions would leave fish with reduced growth, higher stress, and increased vulnerability to predation. Moreover, present-day water depth across the SFE is relatively deeper than historical conditions (Whipple et al., 2012), which may result in more frequent surface access as the rate of gas escape increases (Strand et al., 2005). As such, Delta Smelt metabolic rate may increase with decreasing turbidity. Future research should pursue measuring Delta Smelt SMR and P_crit_ across a range of temperatures and turbidities, and the resulting output could be converted into relevant spatial-temporal models (e.g. metabolic index) to assist in conservation of threatened and endangered species.

The use of respirometry could also be used to quantify Delta Smelt energetic budget and sublethal stress across life stages, which can in turn translate to lower operating costs for FCCL. Recent estimate suggests Delta Smelt costs ∼$84 per fish (Yanagitsuru et al., 2022), which pales in comparison to salmonids that cost between $0.23 to $0.96 per fish (Cavallo et al., 2009; Yanagitsuru et al., 2022). As suggested in this study, it is possible that reduced Delta Smelt anxiety under red lighting (compared to complete darkness or ambient lighting) could save time, improve safety, and reduce operation costs. These improvements are critical as the current US Fish and Wildlife Biological opinion proposes increasing Delta Smelt production target from 33,000 to 125,000 fish (USFWS, 2019), which is a daunting challenge compared to the 55,733 (USFWS, 2022) and 43,725 (USFWS, 2023) fish released in the winter of 2021 and 2022, respectively. In summary, this methodology paves the way for acquiring critical knowledge needed to inform Delta Smelt research and conservation (e.g. the development of metabolic index), and the lessons learned may be applicable to conservation and management of this critically-endangered species by decreasing aquaculture operating costs and improving survival during fish transport and supplementation.

## Methods

### Delta Smelt rearing

These experiments were conducted between October and November 2023 in accordance with protocol no. 23316 in compliance with the Institutional Animal Care and Use Committee at the Center for Aquatic Biology and Aquaculture at University of California Davis.

Delta Smelt were reared at FCCL until 221 days old, transported by truck in black carboys, and placed into rearing tanks (volume = 200 L; n=50 per tank) at the Center for Aquatic Biology and Aquaculture at UC Davis. Temperatures were adjusted at a rate of 1°C every two days until acclimation temperatures were achieved (ranges from 10 to 19°C). Delta Smelt were fed 4% body weight daily (BioPro2 #1 Crum, Bio-Oregon, LongView, WA United States). Delta Smelt rearing tanks had gray walls, and were covered with black (painted) lids made of plastic or Styrofoam. Each cover had a small opening that allowed some ambient light and for food from a belt feeder to fall into the tank during the day. Delta Smelt were acclimated to their respective target temperature for at least two weeks prior to experimentation. Respirometry data was generated with Delta Smelt between 259 and 302 days old (length = 69.7 ± 0.5 mm; weight = 2.6 ± 0.1 g; mean ± SEM).

### Surface restriction behavior

Before dawn and with the assistance of a red-light headlamp, Delta Smelt were gently coaxed from their rearing tank into a 1000 ml container and transferred in a covered bucket into an acclimation tank (20 x 9.5 x 18 cm) to allow the fish to acclimate to the testing area. The testing area is an insulated water bath illuminated overhead with red light, and laterally surrounded by black plastic tarp to remove unintentional stimuli. The temperature was set and kept at their respective acclimation temperature (± 0.5°C) using a water chiller (DBA-150; Arctica, Inglewood, CA, USA) and heatbar (TH-0800S; Finnex, Chicago, IL, USA). After 15 min in the acclimation tank, Delta Smelt were released into one of two testing arenas. The two plastic arenas were completely clear and of the same size (30.5 × 22.5 x 16 cm), and they were stacked on top of each other so that the top arena had surface access whereas the bottom arena did not. Two submersible water pumps constantly delivered aerated water into both arenas. Arenas were rotated 180° after every trial to compensate for any unintentional stimuli. Recordings were continuously captured by a waterproof CCTV camera (LC-F8036; 101AV Inc, Sunnyvale, CA, USA). Fish in the bottom arena were kept 4.5 days without surface access, then moved to the top arena for 2.5 days to monitor their recovery. Fish in the top arena remained there for the entire duration (7 days). Delta Smelt were not fed during the behavioral testing. In total, three trials were conducted: one each at 10, 12, and 19°C.

The angle of the fish was measured using Adobe Illustrator. The *line segment* tool was used to create two lines. One parallel to the water surface that intersects with the snout of the fish, and a second line that connects the snout of the fish to the caudal peduncle of the fish. The angle of the second line was quantified with the *measure tool* in Adobe Illustrator. Due to a tilt in the camera, images were adjusted by 2.2° prior to measurements.

### Predawn respirometry

Delta Smelt respirometry began before dawn between 4 - 6am, and was generally completed between 5 - 7pm. SMR measurements utilized bimodal intermittent respirometry, whereas P_crit_ measurements used classic intermittent respirometry. All respirometry was conducted in a dark room illuminated with overhead red lights (9-31 lumens; 1006 839 413; Ecosmart by Home Depot, Atlanta, USA), and light intensity was measured with a light meter (Extech Light Meter 401025; Industrial Electronics, Inc., Knoxville, TN, USA). The temperature within the water bath continuously maintained within (± 0.3°C) using a heater/chiller (TK-500; Teco, São Paulo, Brazil). O_2_ partial pressure of both the air and the water measured using optical fiber dipping probes (DP-PSt3; PreSens, Regensburg, Germany) inserted into water- and air-tight rubber stoppers and recorded with Autoresp2 or Autoresp3, with each trial having an interval of 90 seconds of flush, 90 seconds of wait, and 600 seconds of measurement. O_2_ levels within the chambers were not allowed to decline below 95% saturation during the SMR measurements to prevent stress.

Respirometry chambers (14 x 6 x 8 cm) were built with clear acrylic to allow visual access to conspecifics, and a clear lid allowed overhead red light to penetrate (Figure 3). Two sets of four chambers were arranged in a row, and all eight chambers were placed in the same water bath. Individual respirometry chambers were fitted with a rubber gasket to prevent leaks, and two perforated acrylic barriers to prevent Delta Smelt from swimming into water inflow or outflow tubing (Figure 3). Each respirometry chamber was equipped with a recirculating pump that passed water over an optical fiber O_2_ probe and returned it to the respirometry chamber (chamber volume ∼28 mL). In addition, the cover of the chamber had a port for a second fiber optic O_2_ probe. Respirometry chambers were placed at the surface of the water bath, with the top portion of the chamber above the water level to produce an air-pocket. Chambers were carefully balanced and parallel with the water surface. As a result, the water table (height of the water within the water bath) kept the water within the respirometry chamber filled to ∼5.7 cm, which is equivalent to a water volume of ∼507 mL and air volume of ∼193 mL during SMR trial. The flush pump (Eheim, Deizisau, Germany) was fitted with a motorized ball valve (USS-MSV00007; Cleveland, OH, USA) to prevent backflow and allow for computer control of flush and measurement periods. Air was gently and continuously bubbled within the water bath, and two water permeable screens were used to limit water and bubble movement that could potentially disturb nearby fish.

Delta Smelt were captured from their rearing tanks before dawn with a 1L beaker and illuminated with a red-light headlamp, then transported to the experimental room in a covered bucket (∼3-minute walk). Delta Smelt were gently coaxed into a plastic tri-corner beaker then gingerly poured into their respective respirometry chambers. The water level within the respirometry chamber was then equalized with the surface level, after which the chamber lid was secured. SMR measurements were generated from 29 cycles (∼6 hours 30 minutes) of respirometry trials.

As stated above, the air pocket within the respirometry chambers must be removed prior to P_crit_ measurements. To minimize disturbance to the fish, the O_2_ probe on the lid of the respirometry chamber was removed to create an opening, after which the flush pump would displace the air pocket with water. Air bubbles sometimes became trapped on the lid, and these were removed with a gentle tilting of the chamber and/or the plugging of the outflow. Once all air bubbles were removed, the O_2_ probe was re-secured to the lid of the respirometry chamber. During P_crit_ measurements, air was replaced with nitrogen bubbling pumped through a ceramic air stone (1DMBDC300; Pentair, Apopka, FL, USA). To keep DO drawdown consistently around -5% per cycle, nitrogen pressure was adjusted to 23 PSI during 100% to 40% DO saturation, 25 PSI during 40 to 20% DO saturation, and 28 PSI during 20% to 0% DO saturation (or until all fish lost equilibrium).

### Respirometry data analysis

Metabolic rates and P_crit_ values were quantified by analyzing the AutoResp™ trial O_2_ Saturation (%O2Sat) output. A running average (window length = 29) was applied to smooth the data (Chabot et al. 2021) prior to conversion to O_2_ concentration. Smoothed O_2_ saturation data were converted to O concentration ([O] mg O L^-1^) using **Equation 1**.

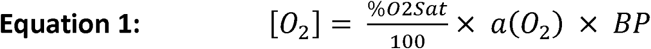

*%O2Sat* is the O_2_ saturation percentage measured via AutoResp, *a(O*_2_) is the temperature-corrected O solubility coefficient (mg O_2_ L^-1^ mm Hg^-1^), and *BP* is the barometric pressure (mmHg). Per-second O_2_ concentration measurements were regressed over time, and the resulting coefficient of this regression (*R*; mg O_2_ L^-1^ s^-1^) was transformed into metabolic rate (MO ; mg O kg^-0.95^ min^-1^) using **Equation 2**.

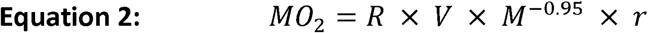

*V* is the volume of the closed respirometer (500 or 700ml depending on whether a trial was quantifying SMR or P respectively), *M* is fish mass (kg^-0.95^) and *r* (60s min^-1^) adjusts the rate to per minute from per second. An allometric scaling exponent of 0.95 was used to account for differences in fish sizes (Lucas et al., 2014).

After the above conversion from O_2_ saturation percentage to O_2_ concentration, SMR measurement periods (n = 29 per fish) were inspected visually. Measurement periods with an R^2^ ≤ 0.70 were individually reviewed, and those indicative of transient temperature fluctuations were discarded (n = 52 of 2,320; 2.2%). Final SMR was calculated using the *calcSMR* function from the *FishMO2* package (Chabot, 2020) incorporating the remaining SMR measurements (n = 24-29 per fish). The remaining measurement periods attributable to the P_CRIT_ trial (n = 12 - 24 per fish) were likewise converted from O_2_ saturation percentage to metabolic rates as described above. The P_CRIT_ was calculated using the *calcO2crit* function from *FishMO2* package. This function uses the metabolic data collected during the P_CRIT_ trial as well as the individual fish’s SMR measurement periods to determine the O_2_ concentration which restricts the metabolic capacity of the fish.

### Statistical analysis to deduce the Influence of fasting, methodology

To assess the impacts of feeding and fasting on MO_2_, Delta Smelt reared at 10 and 12°C were fasted for two days. Predawn respirometry protocols were employed as previously detailed. To assess the impacts of predawn and overnight respirometry protocols, the start time was changed from 4-7am to 3-6pm, with P_crit_ beginning on the following morning between 4 to 7am. To assess the impacts of our inclusion of red light and surface access had on Delta Smelt respirometry, we compared our respirometry results with unpublished 11, 15 and 18°C data collected from juvenile and adult Delta Smelt in 2013 as well as previously published 16°C results from Hammock *et al*. (2017) (only the 0.4 ppt salinity treatment).

Statistical tests were performed using R (version 4.0.3; R Development Core Team, 2013) with packages *nlme* (Pinheiro et al., 2014) and *emmeans* (Lenth, 2021). Statistical differences were analyzed using linear mixed-effects model fitted by restricted maximum likelihood (REML) with *temperature* and another variable of interest (*fed or fasted, predawn or overnight, this study or previous study*) as fixed factors and individual and tank as random factors. Values are reported as mean ± s.e.m., and an alpha of 0.05 was employed for all analyses.

## Acknowledgement

The authors would like to thank Bruce Hammock for his comments on an early manuscript draft. We would also like to thank Department of Water Resource researchers Brittany Davis and Trishelle Tempel for providing context and relevant resources. This work would not be possible without the dedicated efforts by Fish Conservation and Culture Laboratory (FCCL) including Tien Chieh Hung, Troy Alan Stevenson Jr, Joan Lindberg, Luke Ellison, Galen Tigan, and their many staffs.

The latest prototypes for Delta Smelt respirometry research and associated transport, husbandry, and experimental setup were supported by many members of the UC Davis Fangue lab including Sebastian Gonzales, Peter Aronson, Sammuel Huang, Junhan Wang, Kristen Kilaghbian, Kamille Romero, Kylie Tretschok, Mikayla Debarros, and Heather Bell.

Moreover, this research could not have been possible without the help of the many people associated with the earlier prototypes and their fish transport, husbandry, and experimental assistance. We especially thank Felipe La Luz for his contribution to the early respirometry prototypes. We also thank Christine Verhille, Tommy Agosta, Elise

Zarri, Matthias Hasenbein, Lisa Komoroske, Brian Williamson, Oliver Patton, Trinh Nguyen, Noelle Patterson, Jessie Chow, Ryan Young, Izari Chau, Janette Perez Jimenez, Galena Robertson, James Raybould, Colin Turcotte, Bethany Decourten, Jamilynn Poletto, Monica Richmond, Erica Kelly, Mojan Saberi, Jason Dexter, Trinh Nguyen, Krystal Ho, E.P. Scott Weber, Paul Lutes and Erik Hallen, and Frank Loge.

We would like to thank Squidtoons Comic illustrator Dana Song for her Delta Smelt illustration. We are also grateful to the UC Davis Atmospheric Science Department for their long-term measurement and curation of atmospheric barometric data.

## Supplemental Data

Supplemental data has been uploaded to FigShare and can be found at https://figshare.com/s/6ee9ca265ae72a8ece2f

